# Glycolytic enzymes form membrane-less condensates in the malaria parasite *Plasmodium falciparum* by sensing glucose levels

**DOI:** 10.1101/2024.04.08.588540

**Authors:** Ryuta Ishii, Takaya Sakura, Daniel Ken Inaoka, Fuyuki Tokumasu

## Abstract

Recent studies have shown that liquid-liquid phase separation (LLPS) in cells can regulate essential cellular events, including metabolic processes. Glycolytic bodies (G-bodies) are biomolecular condensates formed through the LLPS of glycolytic enzymes, and they accelerate glycolysis to overcome hypoxic stress in several organisms. Although the asexual blood stage (ABS) of the human malaria parasite *Plasmodium falciparum* highly depends on glycolysis for energy production, there have been no reports of the formation of such G-bodies throughout the parasite’s lifecycle. Using fluorescence tagging and live imaging, we found that G-body-like condensates containing phosphofructokinase 9 (PFK9) and phosphoglycerate kinase (PGK) were formed in the parasite cells after long-term culture under conditions of low glucose. These G-body-like structures appeared stable, but membrane staining and osmotic stress experiments suggested that the observed condensates were not associated with a lipid membrane. Notably, PFK9 and PGK formed spatially distinct, non-overlapping condensates, indicating that these structures were formed with different mechanism as compared to the shared, multi-enzyme G-bodies described in other organisms. Whether the condensation of these enzymes enhances or restrains glycolysis under low-glucose stress remains to be determined. These results suggest that the formation of glycolytic enzyme condensates may be an enzyme-specific cellular response that helps the parasite adapt to reductions in blood glucose levels in the host during the ABS.

**Significance statement:** Glycolytic bodies (G-bodies), which are biomolecular condensates formed through the liquid-liquid phase separation of glycolytic enzymes, can accelerate glycolysis to produce energy and overcome hypoxic stress. The parasites that cause malaria depend on glycolysis for energy production, but there have been no reports that these parasites form G-bodies. We demonstrated that membrane-less G-body-like structures formed in media containing low levels of glucose. Two glycolytic enzymes, PFK9 and PGK, formed these condensates, but they did not share a common structure and instead assembled into physically separated condensates, unlike the multi-enzyme G-bodies reported in other organisms. The formation of glycolytic enzyme condensates may be important for the malaria parasite to adapt to reduced blood glucose levels in the host. These results further our understanding of the cellular mechanisms for the survival of malaria parasites.

## Introduction

Malaria is a devastating disease caused by *Plasmodium* parasites transmitted to humans from infected female *Anopheles* mosquitoes. After the asymptomatic liver stage, the parasite releases merozoites into the bloodstream to start the asexual blood stage (ABS), in which the number of parasites increases rapidly up to 10^12^ (1). During these rapid multiplications in the ABS, glycolysis is the parasites’ primary source of energy production, given the fact that several tricarboxylic acid cycle enzymes and the β subunit of adenosine triphosphate (ATP) synthase are dispensable (2-4). Although glycolysis is essential for the blood stage of the parasite, the reaction rates for the glycolysis steps change drastically during intraerythrocytic development (5), which may be difficult to explain by the simple regulation of enzyme expression levels, considering the results of transcriptome analysis (6, 7).

To control the activity of sequential multi-enzyme reactions, cells utilize liquid-liquid phase separation (LLPS) to condense enzymes, substrates, and products within a droplet, which usually allows stable control of the process. For example, purine biosynthesis-related enzymes are compartmentalized in clusters called purinosomes in human cells under purine-depleted conditions to accelerate enzyme reactions (8, 9). Purinosomes contain at least ten enzymes in the purine biosynthesis and salvage pathways, and these enzymes efficiently convert phosphoribosyl pyrophosphate to adenosine monophosphate and guanosine monophosphate (10). Another example of LLPS-induced enzyme reaction activation is the “Glycolytic body (G-body),” which is formed under hypoxia by the condensation of glycolytic enzymes. The assembly of glycolytic enzymes in response to hypoxia was first observed in 2011, when glyceraldehyde-3-phosphate dehydrogenase (GAPDH), the glycolytic enzyme catalyzing the sixth step of glycolysis, was observed to form foci in HeLa cells under hypoxia (11). Subsequently, Miura et al. reported that several glycolytic enzymes were condensed in foci in *Saccharomyces cerevisiae* under hypoxia (12). The yeast Pfk2p has a ‘disordered region’ that lacks a rigid three-dimensional structure. Deletion of this region resulted in failure of the G-body formation, suggesting that Pfk2p served as one of the scaffold proteins of G-bodies, and the disordered region drove the condensation of glycolytic enzymes (13). The detailed components of the G-bodies were identified by isolation and proteome analysis of G-bodies from yeast cells (14). G-bodies contain not only glycolytic enzymes but also chaperones, proteasomal subunits, and fatty acid synthesis-related proteins, suggesting that G-body formation is closely linked with other metabolic pathways and ATP-dependent cellular processes. In these model organisms, glycolysis has been shown to be accelerated by G-body formation to produce more ATP, which is required to increase cell survival (14). Local high concentrations of PFK-1.1, the *Caenorhabditis elegans* ortholog of PFK, were also reported to trigger G-body formation. PFK-1.1 was distributed unevenly in neurons, but helped to satisfy the local energy demands in normoxia (15). G-bodies were thought to be required to narrow down the target area and sustain synaptic function under stress.

Although G-bodies are often regarded as shared, multi-enzyme condensates that improve the efficiency of the glycolytic pathway, the propensity to form higher-order assemblies is not necessarily shared by all glycolytic enzymes, and can differ even between isoforms of the same enzyme. Among mammalian PFK1 isoforms, the liver isoform assembles into filaments in a substrate-dependent manner, whereas the platelet and muscle isoforms do not (16). It has also been shown that glucose flux itself can be sensed through post-translational modification to drive the assembly of glycolytic enzymes, coupling condensate formation to the nutritional state of the cell rather than to hypoxia alone (17). These observations suggest that the composition and physiological role of glycolytic condensates may differ between organisms and conditions, and need to be examined for each system rather than assumed from the yeast model.

Kinetoplastids and diplonemids, such as *Trypanosoma cruzi*, a protozoan parasite causing Chagas disease, have a membrane-bound organelle called a glycosome (18). This structure contains most of the glycolytic enzymes of these protists and may have characteristics similar to those of G-bodies. In *Plasmodium falciparum (Pf)*, LLPS has recently been reported to underlie the assembly of other cellular structures, such as the calcium-inducible condensation of centrins, in which some but not all centrins are condensed together (19). However, there have not been any reports on membrane-bound or membrane-less glycolytic organelle formation in *Apicomplexan* parasites, including *Pf*, although *Pf* highly depends on glycolysis as an energy source.

We hypothesized that G-body-like structures are also formed in the cells of *Pf* parasites and may play a critical role in metabolic regulation. Several of the glycolytic enzymes of *Pf* are disordered proteins, and *Pf*PFK9, the ortholog of yeast Pfk2, has one or more disordered regions, which implies that *Pf*PFK9 may contribute to condensate formation in the parasite cells. In the present study, we investigated condensate formations of glycolytic enzymes, and found that PFK9 and PGK formed stable, membrane-less condensates in the parasite cells under low-glucose conditions, indicating that the processes differ between enzymes rather than a non-specific response of the cytosol. Notably, condensates of PFK9 and PGK did not co-localize but formed spatially distinct structures, suggesting that the processes of these condensates formation in *Pf* are different from the shared, multi-enzyme G-bodies described in other organisms. These results suggest that the condensation of specific glycolytic enzymes may be a unique cellular response that helps the parasite adapt to changes in glucose levels in the human body.

## Results

### PFK9 condensates appeared in *Pf* parasites under low-glucose conditions

To visualize the formation of G-body-like structures in the parasite cells, we generated transgenic parasites that expressed fluorescently tagged PFK9 at the *C*-terminus. The *C*-terminus was selected because tagging of the disordered *N*-terminal sequence may affect the LLPS of PFK9 (Fig. 1A). We used a pKIC-ter plasmid for inducing selection-linked integration (SLI) (20). SLI allows the expression of a target protein to be controlled by an endogenous promoter, which is important for LLPS studies, as slight modifications of the expression level of target proteins easily influence LLPS. The plasmid had a homologous PFK9 C-terminal sequence followed by mNeonGreen (mNG) (21), 2xMyc tag, a tandem sequence of P2A and T2A skip peptides (22), and bacterial neomycin-kanamycin phosphotransferase II (NPTII) as a selection marker. Skip peptides allow the physically separated expression of a tagged target protein and NPTII (Fig. 1B). Diagnostic PCR suggested the successful insertion of the plasmid sequence into the parasite genome (Fig. 1C). The expression of tagged PFK9 was also confirmed by western blotting (Fig. 1D); however, no G-body-like structures were observed in the live imaging (Fig. 1E), unlike a previous report that showed that PFK9 droplet-like structures were formed in the trophozoite state using immunofluorescence assay (23). Therefore, several stimulation methods to facilitate G-body formation were investigated, including heat shock, culture at high parasitemia, and treatment with respiratory chain inhibitors, but none of these conditions induced the formation of G-body-like structures. PFK9 condensates were only observed when the parasites were cultured for more than 72 h in a medium with 5 mM glucose (≈half the glucose concentration of Roswell Park Memorial Institute media (RPMI)) (Fig. 2A and 2B), corresponding to a fasting blood glucose concentration. The clustering percentage, which represents the fraction of pixels with a normalized intensity below a given threshold (24), increased to 123% when the parasite was cultured in a 5 mM low glucose medium (Fig. 2C). Interestingly, there was no drastic difference in transcriptome and metabolome between standard and low-glucose cultured parasites (Fig. S1 and S2). Glucose concentrations less than 5 mM led to early cell death, possibly because of the malfunction of host erythrocytes, and we did not observe any additional PFK9 condensates. These observations suggested that PFK9 condensates were formed through slight but continuous low glucose stress, not acute glucose decreases that do not occur in the human body.

**Figure 1.**
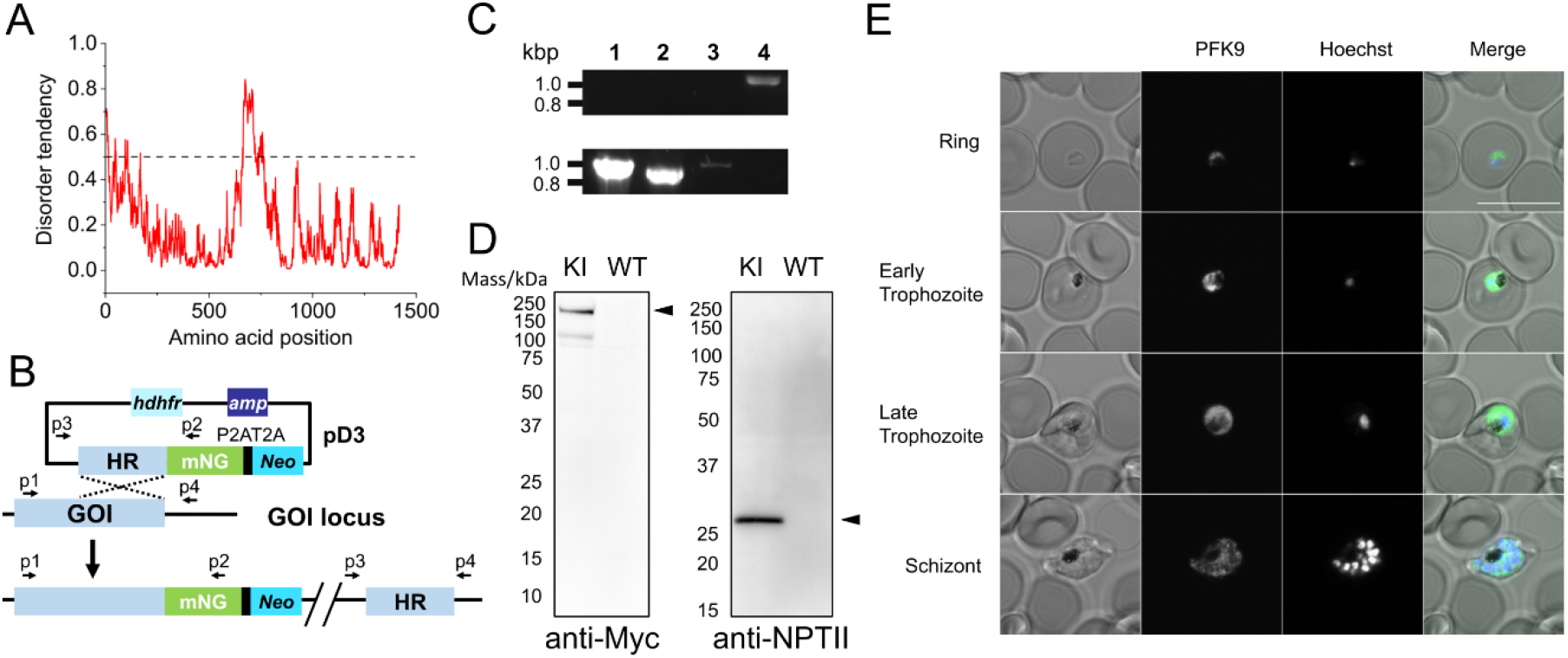
Generation of the *Pf*PFK9-mNG parasite. (A) The result of IUpred2A disorder prediction of *Pf*PFK9 suggesting the existence of *N*-terminal and C-terminal disordered regions. (B) Schematic of pKIC-ter plasmid and SLI. The plasmid has a homologous region of a gene of interest, following mNG, tandem skip peptides P2A-T2A, and NPT II, which shows G418 resistance only when inserted into the parasite genome. The skip peptides induce ribosomal skipping during translation, which allows separate expression of mNG-tagged GOI and NPTII proteins. (C) The results of diagnostic PCR for wild type (WT) and knock-in (KI) strains with different primer sets. Lane 1: p1 + p2, Lane 2: p3 + p4, Lane 3: p2 + p3, Lane 4: p1 + p4. (D) C-Terminal tagging of PFK9 in the *Pf* 3D7 parasite was confirmed by anti-Myc and anti-NPTII western blotting. Application of 2.5 × 10^7^ parasite lysate/lane was used for KI and WT strains. The PFK9-Myc-mNG 190 kDa band and NPTII 29.1 kDa band were only observed in the KI samples, showing successful C-terminal tagging of PFK9. (E) Live imaging of the *Pf*PFK9-mNG parasite. PFK9 was uniformly expressed in the cytoplasm of the parasite throughout the asexual blood stage. G-body-like structures were not observed in the steady state. The scale bar represents 10 µm.

**Figure 2.**
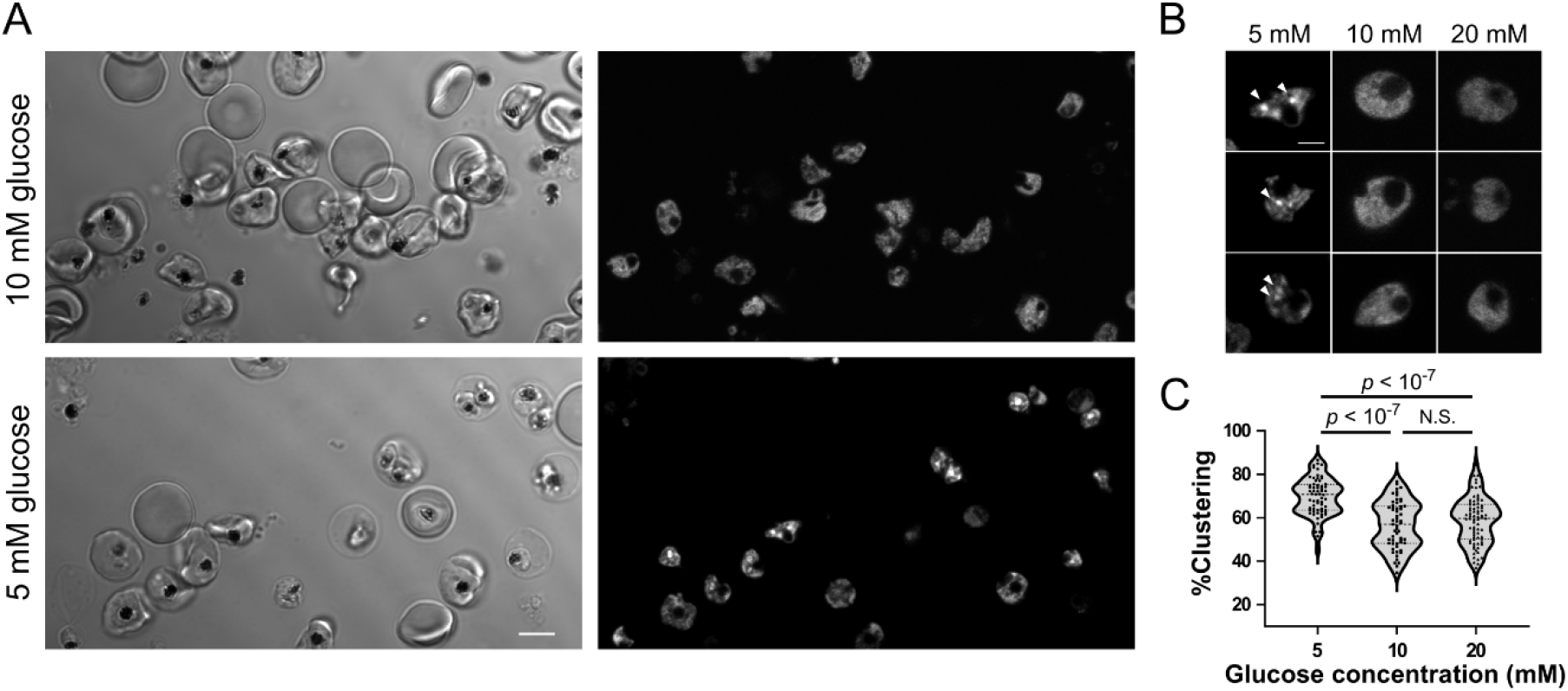
G-body-like structures formed in the *Pf*PFK9-mNG parasite under low glucose conditions. (A) Live imaging of the *Pf*PFK9-mNG parasite in a culture medium containing glucose at the indicated concentrations. The scale bar represents 5 µm. (B) A magnified version of the *Pf*PFK9-mNG parasite cultured in different glucose conditions. The arrowheads indicate the G-body-like PFK9 clusters observed in the 5 mM glucose culture. The scale bar represents 2 µm. (C) Clustering % of PFK9-mNG fluorescence in different glucose concentrations indicated the clustering of PFK9 occurred in a glucose concentration-dependent manner. n = 67 (5 mM), 53 (10 mM), and 59 (20 mM). p values were calculated by Kruskal–Wallis and Steel–Dwass post hoc tests.

### PFK9 condensates behave like membrane-less structures

PFK9 condensates formed after long-term low-glucose culture persisted throughout the observation period. Individual condensates remained detectable and did not fuse with neighboring condensates for at least 30 min (Fig. 3A). The physical properties of PFK9 condensates in parasite cells were examined by fluorescence recovery after photobleaching (FRAP) analysis. However, the small size of condensates with respect to the limited laser spot size did not allow to collect fluorescence signals specifically from condensates because PFK9 signals outside the condensates obscured the signal origins.

**Figure 3.**
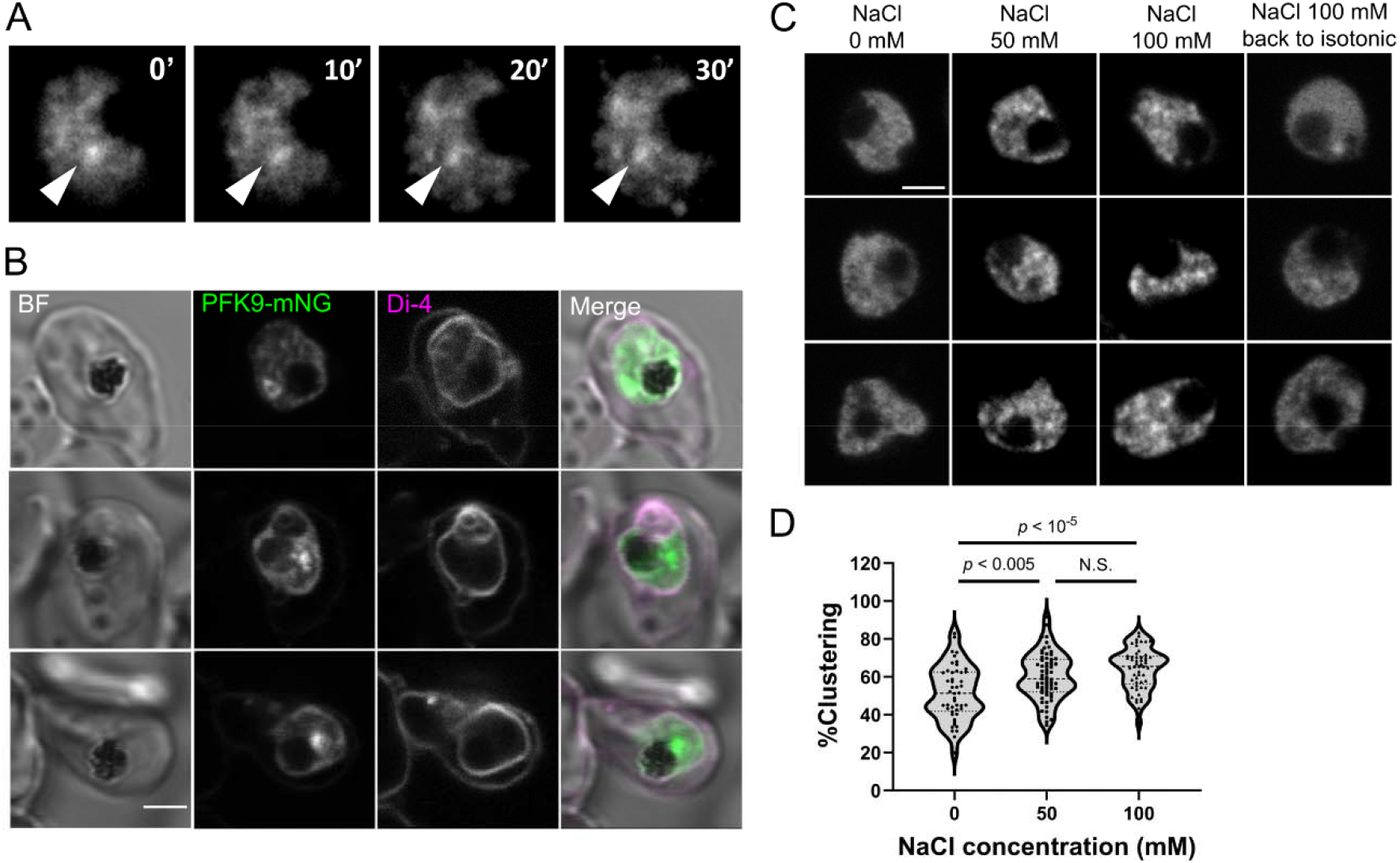
LLPS-like behavior of PFK9 clusters. (A) Time-lapse images of the *Pf*PFK9-mNG parasite over 30 min. The arrowheads indicate a representative stable G-body-like PFK9 cluster. (B) The *Pf*PFK9-mNG parasite was cultured in a low glucose medium and stained with the membrane probe Di-4-ANEPPDHQ for 30 min. The parasite plasma membrane and some inner membranes showed fluorescence, whereas the G-body-like PFK9 clusters were not stained with the dye. The scale bar represents 2 µm. (C) Live imaging of *Pf*PFK9-mNG parasites suspended in CM with the addition of the indicated NaCl concentration. PFK9 clusters were observed in the parasite cytosol and disappeared when the CM was reverted to isotonic conditions. The scale bar represents 2 µm. (D) Clustering % of the *Pf*PFK9-mNG parasite indicated the clustering of PFK9 under osmotic stress. n = 48 (0 mM), 63 (50 mM), and 53 (100 mM). p values were calculated by Kruskal–Wallis and Steel–Dwass post hoc tests.

To investigate whether the PFK9 condensates were membranous structures, parasitized erythrocytes were stained with Di-4-ANEPPDHQ, a dye that shows fluorescence only when in a phospholipid membrane (25). The G-body-like condensates were devoid of the Di-4-ANEPPDHQ signal, whereas the parasite membranes showed fluorescence, indicating that the G-body-like condensates were membrane-less organizations formed through LLPS (Fig. 3B).

For further characterization of the PFK9 condensates, hyperosmotic experiments were performed to study condensate formation with temporal changes in cellular protein concentrations. Recent studies have shown that hyperosmotic stress can cause LLPS of several proteins because osmotic pressure compresses the cell volume and increases the protein concentration and macromolecular crowding (26). In addition, the local concentration of PFK was likely the key determinant of G-body formation in *C. elegans* (27). After the induction of hyperosmotic stress by the addition of 50 mM of sodium chloride, the rapid formation of PFK9 condensates was observed within 5 min (Fig. 3C and 3D). These structures dissolved within 5 min after the medium was returned to isotonic conditions (Fig. 3C), whereas the condensates formed under long-term low glucose persisted throughout the 30 min observation (Fig. 3A). These observations suggested that an increase in the local PFK9 concentration and macromolecular crowding could induce the formation of PFK9 conden t sates in the parasite.

### PFK9 and PGK form spatially distinct condensates under low glucose conditions

To investigate whether condensate formation also occurred for other glycolytic enzymes, several transgenic parasites expressing mNG-tagged glycolytic enzymes were generated using SLI. Of the nine glycolytic enzymes, six enzymes, in addition to PFK9, were successfully tagged and had the expression of the tagged proteins confirmed (Fig. S3). However, the tagging of fructose-bisphosphate aldolase (FBPA; PF3D7_1444800), GAPDH (PF3D7_1462800), and enolase (ENO; PF3D7_1015900) was not successful, possibly because of the toxic effects of the *C*-terminus tagging of these genes, and further investigation is needed. The obtained transgenic parasites were cultured under low glucose conditions for 80 h, and only PGK (PF3D7_0922500) formed similar condensates to that of PFK9 in the cytosol (Fig. 4). These results indicated that condensate formation under low glucose conditions was restricted to specific enzymes, PFK9 and PGK, rather than reflecting a stress-induced non-specific aggregation of the cytosol.

**Figure 4.**
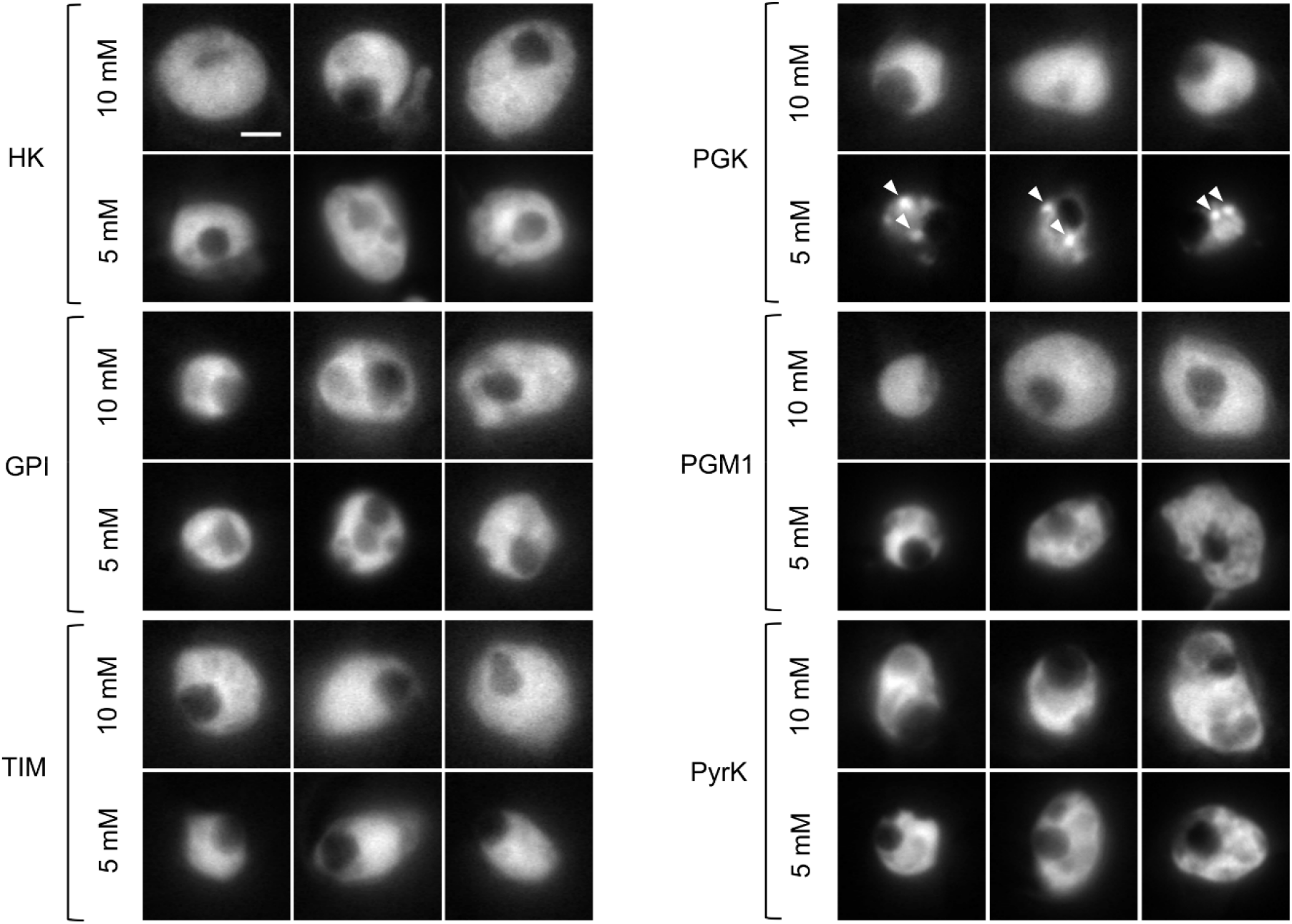
PGK also formed G-body-like clusters under low glucose conditions. Representative live images of the parasites that expressed tagged glycolytic enzymes cultured in different glucose concentrations for 80 h. Only PGK formed clusters in 5 mM glucose culture, as indicated by the arrowheads. The scale bar represents 2 µm. Enzyme abbreviations: HK, hexokinase; GPI, glucose-6-phosphate isomerase; PFK9, phosphofructokinase 9; FBPA, fructose-bisphosphate aldolase; TIM, triosephosphate isomerase; GAPDH, glyceraldehyde-3-phosphate dehydrogenase; PGK, phosphoglycerate kinase; PGM1, phosphoglycerate mutase 1; ENO, enolase; PyrK, pyruvate kinase.

To test whether PFK9 and PGK are recruited into the same condensates, as would be expected for a shared, multi-enzyme G-body, we generated a dual-tagged line co-expressing differentially labeled PFK9 and PGK and imaged it under low glucose conditions (Fig. 5A). Unexpectedly, PFK9 and PGK localized to physically separate, non-overlapping foci rather than to common condensates (Fig. 5B). Thus, although both enzymes condense in response to the same stimulus, they form distinct, spatially segregated condensates.

**Figure 5.**
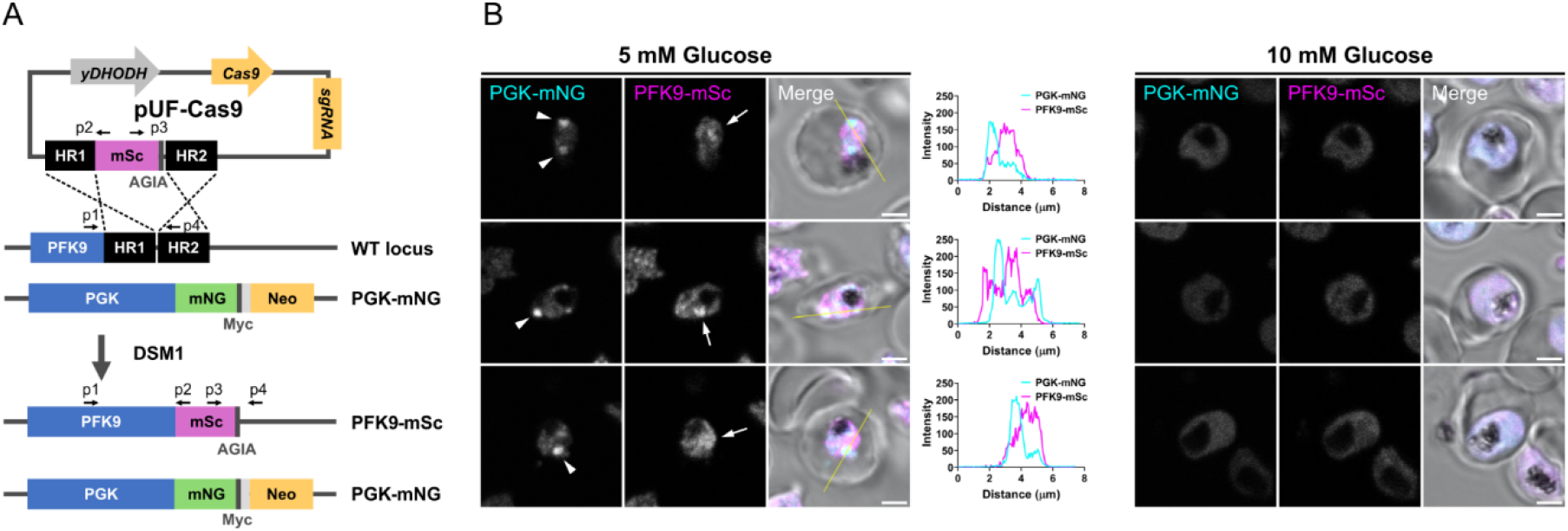
PFK9 and PGK formed spatially distinct condensates under low glucose conditions. (A) Schematic of the pUF-Cas9 plasmid and the strategy for generating the double knock-in parasite. The plasmid has two homologous regions of PFK9 (HR1 and HR2) flanking mScarlet (mSc) and an AGIA tag, together with a Cas9 expression cassette, an sgRNA, and yDHODH, which confers DSM1 resistance. C-terminal tagging of PFK9 with mSc was performed in the PGK-mNG parasite, resulting in a parasite expressing both PFK9-mSc and PGK-mNG. p1–p4 indicate the primers used for diagnostic PCR. (B) Representative live images of the double knock-in parasite cultured in a culture medium containing glucose at the indicated concentrations for 80 h. The arrowheads and arrows indicate the PGK-mNG and PFK9-mSc condensates, respectively. Intensity profiles of the PGK-mNG and PFK9-mSc along the yellow lines shown in the merged images. Both enzymes formed condensates in the parasite cytosol in the 5 mM glucose culture, but the two signals did not overlap at the resolution of confocal microscopy. The scale bar represents 2 µm.

## Discussion

In the present study, PFK9 was the primary enzyme investigated because 1) glycolysis is one of the most important processes in the *Pf* lifecycle; 2) PFK9 is the most investigated protein of the G-body components; and 3) *Pf*PFK9 also has disordered regions, which are required for G-body formation in yeast. It was unexpected that PFK9 did not form condensates in a steady state, probably because the parasite was already in a hypoxic condition (30), and the function of G-bodies in *Pf* may differ from that in other organisms. In addition, it should be noted that RPMI 1640-based standard culture medium provides for a higher growth rate than human serum because of the supraphysiological nutrient levels in RPMI (31, 32), and the metabolic conditions in cultured parasites and the need for G-bodies might differ from those of the parasite in vivo. This perspective led to the successful observation of condensates under fasting blood sugar-level conditions.

The long-term low glucose culture results for the seven parasites that expressed tagged glycolytic enzymes suggested that PFK9 and PGK each formed their own condensates under low glucose conditions. In the present study, C-terminal-tagged FBPA, GAPDH, and ENO-expressing parasites were not obtained successfully. The crystal structure analyses of FBPA and GAPDH have suggested that the working units are a homodimer and a homotetramer, respectively (33, 34). The results of recombinant *Pf*ENO purification also indicated that a homodimer was formed (35); therefore, the C-terminal tagging of these proteins may have disrupted multimer formation and resulted in the cell death of the parasite. Crystal structure analysis of PGK demonstrated that it did not have a disordered region (36), suggesting that it is unlikely to drive homotypic phase separation on its own, and its condensates are more likely to be heterotypic, with PGK recruited as a client to a distinct scaffold, such as another disordered protein or RNA, rather than sharing a condensate with PFK9. In this view, the two enzymes may simply belong to different scaffold– client systems. Their formation of separate condensates therefore does not exclude the possibility that the disordered region of PFK9 acts as a scaffold for its own condensate (37). Consistent with a disorder-driven scaffolding of PFK9, the purified disordered regions of PFK9 with and without a GFP tag aggregated even at low concentrations (< 2 µM), suggesting strong intermolecular or intramolecular interactions of the PFK9 occurred in these regions (Fig. S4).

A notable feature of the Pf condensates is that PFK9 and PGK did not co-localize but occupied physically separate, non-overlapping condensates. This organization differs from the canonical G-body, in which sequential glycolytic enzymes co-condense so that intermediates can be channeled directly between active sites to accelerate flux. Because PFK9 and PGK bracket the boundary between the ATP-investment and ATP-payoff phases of glycolysis, their independent condensation may represent separate regulatory responses rather than the assembly of shared metabolons. The functional advantage or disadvantage of this arrangement is not yet resolved and could plausibly act in either direction. On one hand, local condensate of each enzyme could enhance the corresponding reaction and, by extension, overall flux under low-substrate conditions. On the other hand, the physical separation of the two enzymes could instead sequester PFK9 and PGK away from the bulk cytosol, dampening rather than accelerating glycolysis. The metabolome data do not provide a suggestion to this question: the levels of glycolytic intermediates and adenylate pools were unchanged between low- and standard-glucose conditions (Fig. S2 and Table S1), which is consistent both with an absence of flux enhancement and with maintained or elevated activity that keeps metabolite pools constant despite reduced extracellular glucose. Distinguishing between these possibilities will require direct measurements of glycolytic flux and parasite growth under matched conditions.

The results for membrane staining and osmotic stress conditions suggested that the condensation of PFK9 was membrane-less. Interestingly, unlike condensates formed after long-term low glucose culture, which showed no detectable change over the observation period, the osmotically induced condensates dissolved as soon as the stress was removed. The difference between the two conditions may indicate that the physical properties of PFK9 condensates depend on how they are induced, although our data do not allow us to distinguish the underlying mechanisms. The exact mechanism for triggering condensate formation was not identified in the present study. However, considering that the PFK9 expression level did not change even after low glucose culture (Fig. S1), there is likely to be some machinery for increasing the local PFK9 interactions and concentrations, such as post-translational modifications or an increase in other cluster-facilitating proteins. Another possibility for a PFK9-condensation facilitator is a non-protein scaffold molecule, such as RNA. It has been reported that long noncoding RNA could assemble a metabolon of several glycolytic enzymes to overcome metabolic stress (40), and a similar phenomenon may occur in the parasite cytosol. Fusing an RNase to the *C*-terminus of PFK9 may be able to determine the role of RNA in condensate formation, as has been performed in yeast (41). A limitation of the present study was the lack of direct observations of liquid-like behaviors in PFK9 condensates because of technical difficulties with the FRAP analysis and the difficulty of observing the fusion of PFK9 condensates in small parasite bodies. Further investigations are required to investigate the LLPS of PFK9 using other methods, such as observations using fluorescence correlation spectroscopy, which detects changes in molecular diffusion when condensates fuse with each other and become larger clusters (42).

Because 5 mM glucose corresponds to a fasting blood sugar level, the formation of PFK9 and PGK condensates under sustained low-glucose culture may represent a cellular response by which the parasite adapts to prolonged reductions in host blood glucose. The rate at which the parasite consumes its primary energy source may be tuned by the LLPS of these glycolytic enzymes in accordance with the nutritional condition of the host, which may differ appreciably after an increase in the parasite load (43).

More studies will be needed to clarify the significance and nature of PFK9 and PGK condensation by measuring the growth rates and glycolytic activity of the parasite under physiological conditions. However, given the fact that the metabolite levels of glycolysis did not change even after long-term low glucose stress (Fig. S2 and Table S1), PFK9 condensation may contribute to maintaining glycolysis activity under conditions of low substrate levels.

In summary, we found that PFK9 and PGK form G-body-like membrane-less condensates in low glucose conditions. Rather than assembling into a single shared structure, PFK9 and PGK formed enzyme-specific, spatially distinct condensates, indicating that these *Pf* structures diverge from the canonical multi-enzyme G-bodies of other organisms. Whether this arrangement enhances or restrains glycolysis under low-glucose stress remains to be determined.

## Materials and methods

### Culture of *Pf* parasite

*Pf* 3D7 parasites (44) were maintained in complete medium [CM; RPMI 1640 medium with 25 mM HEPES, supplemented with 25 mg/L gentamicin, 50 mg/L hypoxanthine, 23.8 mM sodium bicarbonate, and 0.5% Albumax II at 2% hematocrit of O+ human erythrocytes at 37 °C with a mixed gas (5% O_2_, 5% CO_2_, and 90% N_2_)]. The parasite culture was diluted every 2 or 3 days to maintain the parasitemia between 0.1% and 5%, typically.

### Plasmid construction

For the fluorescent tagging of glycolytic enzymes, SLI was induced by transfection of a pKIC-ter plasmid derived from a pD3 plasmid (45, 46), which has human dihydrofolate reductase and NPTII as selection markers against WR99210 and G418, respectively. Myc tag, a green fluorescent protein mNG tag, and a skip peptide P2A and T2A tandem sequence were synthesized and cloned into pD3 by a seamless ligation cloning extract (SLiCE) reaction (47), resulting in the plasmid pKIC-ter. The pKIC-ter plasmid was restricted with NotI, and the C-terminal sequence of the gene of interest amplified from the 3D7 genomic DNA (gDNA) or synthesized DNA (Table S2) was inserted using SLiCE. The plasmid for CRISPR/Cas9-mediated knock-in was constructed using pUF-Cas9-pre-sgRNA (a gift from Josh Beck; Addgene plasmid #137845; RRID: Addgene_137845). The single guide RNA (sgRNA) targeting the C-terminal region of PFK9 was inserted into BtgZI site of pUF-Cas9-pre-sgRNA, resulting in the pUF-Cas9-PFK9-sgRNA plasmid. Subsequently, a fragment containing two homology regions of PFK9 and mScarlet with an AGIA tag, generated by overlap PCR, was inserted into NcoI site of the pUF-Cas9-PFK9-sgRNA plasmid (48). The primers used for the amplification and the sgRNA are summarized in Table S3.

### Transfection and drug selection of *Pf*

Erythrocytes were suspended in cytomix (49) [120 mM KCl, 0.15 mM CaCl_2_, 10 mM K_2_HPO_4_/KH_2_PO_4_ (pH 7.6), 25 mM HEPES-KOH (pH 7.6), 2 mM ethylene glycol tetraacetic acid (pH 7.6), and 5 mM MgCl_2_, adjusted to pH 7.6 with KOH], centrifuged, and the supernatant was removed. Fifty micrograms of plasmid DNA and 150 µL of packed erythrocytes were mixed, and the volume was made up to 400 µL with cytomix in a 0.2 cm electroporation cuvette. Electroporation was performed with GenePulser (Bio-Rad Laboratories, Hercules, CA, USA) using the conditions: voltage = 310 V; capacitance = 975 µF; resistance = ∞; and cuvette = 2 mm. Erythrocytes were washed with cold RPMI twice and then suspended in CM. The parent parasites were inoculated to the electroporated erythrocytes at 0.1% – 0.3% parasitemia and cultured in a flask as described above. When parasitemia reached ≈5%, drug selection was initiated by adding final 1 nM WR99210 or 1 μM DSM1 to the culture, and culturing was continued for 10 days. For the SLI strategy, following parasite re-emergence, 400 µg/mL G418 was added to the culture for 10 days to select parasites with genome insertion of the plasmid sequence.

### Diagnostic PCR

The gDNA of the parasite was purified with DNeasy Blood & Tissue kits (QIAGEN, Hilden, Germany). Insertion of the plasmid sequences was confirmed by PCR using gDNA as templates. Primers p1 to p4 (Table S3) were used for genotyping.

### Western blotting

The parasite culture was pelleted and suspended in PBS containing 1 × complete protease inhibitor cocktail (Roche, Basel, Switzerland). Saponin (at a final concentration of 0.15%) was added to the solution and incubated on ice for 8 min. The parasite was pelleted by centrifugation at 16,900 × *g* and washed twice with PBS containing 1 × complete protease inhibitor cocktail. NuPAGE LDS sample buffer containing NuPAGE sample reducing agent (Invitrogen, Waltham, MA, USA) was added to the pellet to obtain 10^6^ parasites/µL and heated at 70 °C for 10 min. A sample of the prepared solution (10 to 25 µL) was applied to a NuPAGE 4% to 12% Bis-Tris mini protein gradient gel, and SDS-PAGE was performed in MOPS SDS running buffer at 200 V for 40 min. Membrane transfer to the polyvinylidene difluoride membrane was performed using the XCell II blot module and NuPAGE transfer buffer at 30 V for 1 h. The membrane blocking was performed with Bullet Blocking One (Nacalai, Kyoto, Japan) for 5 min at room temperature. PBS containing 5% Bullet Blocking One was used for antibody dilution. The antibodies and reaction conditions used in the present study are summarized in Table S4. SuperSignal West Pico PLUS chemiluminescent substrate (Invitrogen) was used to generate chemiluminescent signals, and images were taken using a Luminograph II (Atto, Tokyo, Japan) or FUSION-FX6.EDGE (Vilber, Collégien, France).

### G-body induction

Low glucose CM [glucose-free RPMI 1640 (Thermo Fisher Scientific, Waltham, MA, USA) medium supplemented with 25 mg/L gentamicin, 50 mg/L hypoxanthine, 25 mM HEPES, 23.8 mM sodium bicarbonate, 0.5% Albumax II, and 5 mM glucose] was used for the induction of G-body formation. The parasite cultured in the same CM but supplemented with 10 mM glucose was used as a negative control. The 3D7 parasite expressing mNG-tagged glycolytic enzymes was diluted in 2% hematocrit low glucose CM at 0.1%–0.3% parasitemia and maintained for 3 to 4 days until G-bodies were observed by microscopy.

### Live imaging of the parasite

Hoechst 33258 was added to the parasite culture at 2 µg/mL for nuclear staining. After 30 min of incubation at 37 °C, the parasite culture was washed with CM, centrifuged, applied on a microscope slide, and sealed with a coverslip. Images were taken by a LSM780 (Zeiss, Oberkochen, Germany) or DMi8 (Leica, Wetzlar, Germany) camera.

### Calculation of clustering percentage

The image analyses were performed using ImageJ Fiji (50). For calculating the clustering percentage (24), parasite cell images were automatically cropped after performing ‘subtract background,’ thresholding using Huang’s fuzzy thresholding method, running ‘analyze particle’ and ‘enhance contrast.’ The clustering percentage was calculated from the areas measured after thresholding at an intensity of 90 divided by the areas measured without thresholding. Graphs were prepared using GraphPad Prism (GraphPad Software, La Jolla, MA, USA), and statistical analyses were performed with EZR (Saitama Medical Center, Jichi Medical University, Saitama, Japan) (51).

### Lipid membrane staining

The PFK9-mNG expressing parasite was cultured in 10 or 5 mM glucose-containing CM for 4 days. The parasite was centrifuged, and the supernatant was replaced with 5 mM glucose-containing RPMI 1640 with Di-4-ANEPPDHQ (Thermo Fisher Scientific) at 2 µg/mL, and the mixture was incubated at 37 °C for 30 min. The parasite was washed with 10 or 5 mM glucose-containing CM, centrifuged again, applied on a microscope slide, and sealed with a coverslip. Images were taken using an LSM780 microscope.

### Osmotic stress induction

The PFK9-mNG expressing parasite cultured in the standard CM was centrifuged and suspended in a medium containing 50 or 100 mM sodium chloride. After 5 min of incubation at 37 °C, the parasite was centrifuged, applied on a microscope slide, and sealed with a coverslip. Images were taken using a LSM780 microscope. Graphs were prepared using GraphPad Prism, and statistical analyses were performed with EZR.

### Metabolome analysis

The PFK9-mNG expressing parasite was purified by magnetic separation using a MACS separator with LS columns (Miltenyi Biotech, Bergisch Gladbach, Germany) and inoculated to CM containing erythrocytes at 2.5% hematocrit. After 6 h of incubation at 37 °C, sorbitol synchronization was performed to obtain the ring stage parasites (6 h synchronization window), which were suspended in 10 or 5 mM glucose-containing CM, and incubated at 37 °C for 78–80 h. The parasite was purified by magnetic separation, centrifuged, and suspended in methanol, followed by freezing in liquid nitrogen. The quantification of 116 metabolites and statistical analysis was performed by Human Metabolome Technologies.

### Transcriptome analysis

The PFK9-mNG expressing parasite was purified by magnetic separation and inoculated to CM containing erythrocytes at 2.5% hematocrit. After 3 h of incubation at 37 °C, sorbitol synchronization was performed to obtain the ring stage parasites (3 h synchronization window), which were suspended in 10 or 5 mM glucose-containing CM, and incubated at 37 °C for 24 or 36 h. The total RNA was purified with a PureLink RNA Mini kit (Thermo Fisher Scientific) and sent to Novogene for reverse transcription, sequencing, mapping to the *Pf* 3D7 genome, and statistical analysis. The data have been deposited with links to BioProject accession number PRJDB17842 in the DDBJ BioProject database (https://www.ddbj.nig.ac.jp/bioproject/).

## Supporting information

TableS1_Metabolome_Quant

## Acknowledgments and funding sources

This work was supported by JSPS KAKENHI Grant Number JP22K07045 and JP25K10334. This work was also partially funded by the Nagasaki University WISE Programme. We thank Eizo Takashima and Kazuhide Yahata for providing anti-EXP2 and anti-AGIA antibodies, respectively. We thank Masaharu Eriguchi for support in generating and characterizing of the transgenic parasite strains. The Japanese Red Cross Society provided human erythrocytes. This study was conducted (in part) at the Joint Usage / Research Center on Tropical Disease, Institute of Tropical Diseases, Nagasaki University, Japan. Victoria Muir, PhD, from Edanz (https://jp.edanz.com/ac) edited a draft of this manuscript.

## References

1. Duffey M, et al. (2021) Assessing risks of Plasmodium falciparum resistance to select next-generation antimalarials. Trends in Parasitology 37(8):709–721.

2. Salcedo-Sora JE, Caamano-Gutierrez E, Ward SA, & Biagini GA (2014) The proliferating cell hypothesis: a metabolic framework for Plasmodium growth and development. Trends Parasitol 30(4):170–175.

3. Evers F, et al. (2021) Composition and stage dynamics of mitochondrial complexes in Plasmodium falciparum. Nature Communications 12(1):3820.

4. Sturm A, Mollard V, Cozijnsen A, Goodman CD, & McFadden GI (2015) Mitochondrial ATP synthase is dispensable in blood-stage Plasmodium berghei rodent malaria but essential in the mosquito phase. Proc Natl Acad Sci U S A 112(33):10216–10223.

5. Shivapurkar R, et al. (2018) Evaluating antimalarial efficacy by tracking glycolysis in Plasmodium falciparum using NMR spectroscopy. Sci Rep 8(1):18076.

6. Chappell L, et al. (2020) Refining the transcriptome of the human malaria parasite Plasmodium falciparum using amplification-free RNA-seq. BMC Genomics 21(1):395.

7. Kucharski M, et al. (2020) A comprehensive RNA handling and transcriptomics guide for high-throughput processing of Plasmodium blood-stage samples. Malar J 19(1):363.

8. An S, Kumar R, Sheets ED, & Benkovic SJ (2008) Reversible Compartmentalization of de Novo Purine Biosynthetic Complexes in Living Cells. Science 320(5872):103–106.

9. Zhao H, et al. (2015) Quantitative analysis of purine nucleotides indicates that purinosomes increase de novo purine biosynthesis. J Biol Chem 290(11):6705–6713.

10. Pedley AM, Pareek V, & Benkovic SJ (2022) The Purinosome: A Case Study for a Mammalian Metabolon. Annu Rev Biochem 91:89–106.

11. Agbor TA, et al. (2011) Small ubiquitin-related modifier (SUMO)-1 promotes glycolysis in hypoxia. J Biol Chem 286(6):4718–4726.

12. Miura N, et al. (2013) Spatial reorganization of Saccharomyces cerevisiae enolase to alter carbon metabolism under hypoxia. Eukaryot Cell 12(8):1106–1119.

13. Fuller GG, et al. (2020) RNA promotes phase separation of glycolysis enzymes into yeast G bodies in hypoxia. eLife 9:e48480.

14. Jin M, et al. (2017) Glycolytic Enzymes Coalesce in G Bodies under Hypoxic Stress. Cell Rep 20(4):895–908.

15. Jang S, et al. (2016) Glycolytic Enzymes Localize to Synapses under Energy Stress to Support Synaptic Function. Neuron 90(2):278–291.

16. Webb BA, Dosey AM, Wittmann T, Kollman JM, & Barber DL (2017) The glycolytic enzyme phosphofructokinase-1 assembles into filaments. J Cell Biol 216(8):2305–2313.

17. Wang H, et al. (2024) Organization of a functional glycolytic metabolon on mitochondria for metabolic efficiency. Nat Metab 6(9):1712–1735.

18. Quiñones W, et al. (2020) Structure, Properties, and Function of Glycosomes in Trypanosoma cruzi. Frontiers in Cellular and Infection Microbiology 10.

19. Voß Y, Klaus S, Lichti NP, Ganter M, & Guizetti J (2023) Malaria parasite centrins can assemble by Ca2+-inducible condensation. PLOS Pathogens 19(12):e1011899.

20. Spielmann T, et al. (2017) Selection linked integration (SLI) for endogenous gene tagging and knock sideways in Plasmodium falciparum parasites. Protocol Exchange.

21. Shaner NC, et al. (2013) A bright monomeric green fluorescent protein derived from Branchiostoma lanceolatum. Nature Methods 10(5):407–409.

22. Liu Z, et al. (2017) Systematic comparison of 2A peptides for cloning multi-genes in a polycistronic vector. Sci Rep 7(1):2193.

23. Mony BM, Mehta M, Jarori GK, & Sharma S (2009) Plant-like phosphofructokinase from Plasmodium falciparum belongs to a novel class of ATP-dependent enzymes. International Journal for Parasitology 39(13):1441–1453.

24. Ladouceur A-M, et al. (2020) Clusters of bacterial RNA polymerase are biomolecular condensates that assemble through liquid–liquid phase separation. Proceedings of the National Academy of Sciences 117(31):18540–18549.

25. Jin L, Millard AC, Wuskell JP, Clark HA, & Loew LM (2005) Cholesterol-enriched lipid domains can be visualized by di-4-ANEPPDHQ with linear and nonlinear optics. Biophys J 89(1):L04–06.

26. Jalihal AP, et al. (2021) Hyperosmotic phase separation: Condensates beyond inclusions, granules and organelles. Journal of Biological Chemistry 296.

27. Jang S, et al. (2021) Phosphofructokinase relocalizes into subcellular compartments with liquid-like properties in vivo. Biophys J 120(7):1170–1186.

28. Tokumasu F, et al. (2005) Band 3 modifications in Plasmodium falciparum-infected AA and CC erythrocytes assayed by autocorrelation analysis using quantum dots. Journal of Cell Science 118(5):1091–1098.

29. Hwang J, Gheber LA, Margolis L, & Edidin M (1998) Domains in Cell Plasma Membranes Investigated by Near-Field Scanning Optical Microscopy. Biophysical Journal 74(5):2184–2190.

30. Branco A, Francisco D, & Hanscheid T (2018) Is There a ‘Normal’ Oxygen Concentration for in vitro Plasmodium Cultures? Trends Parasitol 34(10):811–812.

31. Pillai AD, et al. (2012) Solute restriction reveals an essential role for clag3-associated channels in malaria parasite nutrient acquisition. Mol Pharmacol 82(6):1104–1114.

32. Desai SA (2013) Insights gained from P. falciparum cultivation in modified media. ScientificWorldJournal 2013:363505.

33. Kim H, Certa U, Döbeli H, Jakob P, & Hol WGJ (1998) Crystal Structure of Fructose-1,6-bisphosphate Aldolase from the Human Malaria Parasite Plasmodium falciparum. Biochemistry 37(13):4388–4396.

34. Robien MA, et al. (2006) Crystal structure of glyceraldehyde-3-phosphate dehydrogenase from Plasmodium falciparum at 2.25 Å resolution reveals intriguing extra electron density in the active site. Proteins: Structure, Function, and Bioinformatics 62(3):570–577.

35. Pal-Bhowmick I, et al. (2004) Cloning, over-expression, purification and characterization of Plasmodium falciparum enolase. European Journal of Biochemistry 271(23-24):4845–4854.

36. Smith CD, Chattopadhyay D, & Pal B (2011) Crystal structure of Plasmodium falciparum phosphoglycerate kinase: Evidence for anion binding in the basic patch. Biochemical and Biophysical Research Communications 412(2):203–206.

37. Banani SF, et al. (2016) Compositional Control of Phase-Separated Cellular Bodies. Cell 166(3):651–663.

38. Patel A, et al. (2017) ATP as a biological hydrotrope. Science 356(6339):753–756.

39. McSwiggen DT, Mir M, Darzacq X, & Tjian R (2019) Evaluating phase separation in live cells: diagnosis, caveats, and functional consequences. Genes Dev 33(23-24):1619–1634.

40. Zhu Y, et al. (2022) The long noncoding RNA glycoLINC assembles a lower glycolytic metabolon to promote glycolysis. Molecular Cell 82(3):542-554.e546.

41. Fuller GG, et al. (2020) RNA promotes phase separation of glycolysis enzymes into yeast G bodies in hypoxia. Elife 9.

42. Yu O & Masataka K (2009) Multipoint fluorescence correlation spectroscopy with total internal reflection fluorescence microscope. Journal of Biomedical Optics 14(1):014030.

43. Binh TQ, et al. (1997) Glucose metabolism in severe malaria: Minimal model analysis of the intravenous glucose tolerance test incorporating a stable glucose label. Metabolism 46(12):1435–1440.

44. Walliker D, et al. (1987) Genetic analysis of the human malaria parasite Plasmodium falciparum. Science 236(4809):1661–1666.

45. Riglar DT, et al. (2011) Super-resolution dissection of coordinated events during malaria parasite invasion of the human erythrocyte. Cell Host Microbe 9(1):9–20.

46. Morita M, et al. (2018) PV1, a novel Plasmodium falciparum merozoite dense granule protein, interacts with exported protein in infected erythrocytes. Sci Rep 8(1):3696.

47. Motohashi K (2015) A simple and efficient seamless DNA cloning method using SLiCE from Escherichia coli laboratory strains and its application to SLiP site-directed mutagenesis. BMC Biotechnology 15(1):47.

48. Morita M, et al. (2021) AGIA Tag System for Ultrastructural Protein Localization Analysis in Blood-Stage Plasmodium falciparum. Front Cell Infect Microbiol 11:777291.

49. van den Hoff MJ, Moorman AF, & Lamers WH (1992) Electroporation in ‘intracellular’ buffer increases cell survival. Nucleic Acids Res 20(11):2902.

50. Schindelin J, et al. (2012) Fiji: an open-source platform for biological-image analysis. Nat Methods 9(7):676–682.

51. Kanda Y (2013) Investigation of the freely available easy-to-use software ‘EZR’ for medical statistics. Bone Marrow Transplant 48(3):452–458.

