## Supplementary Materials for "Glycolytic enzymes form membrane-less condensates in the malaria parasite *Plasmodium falciparum* by sensing glucose levels"

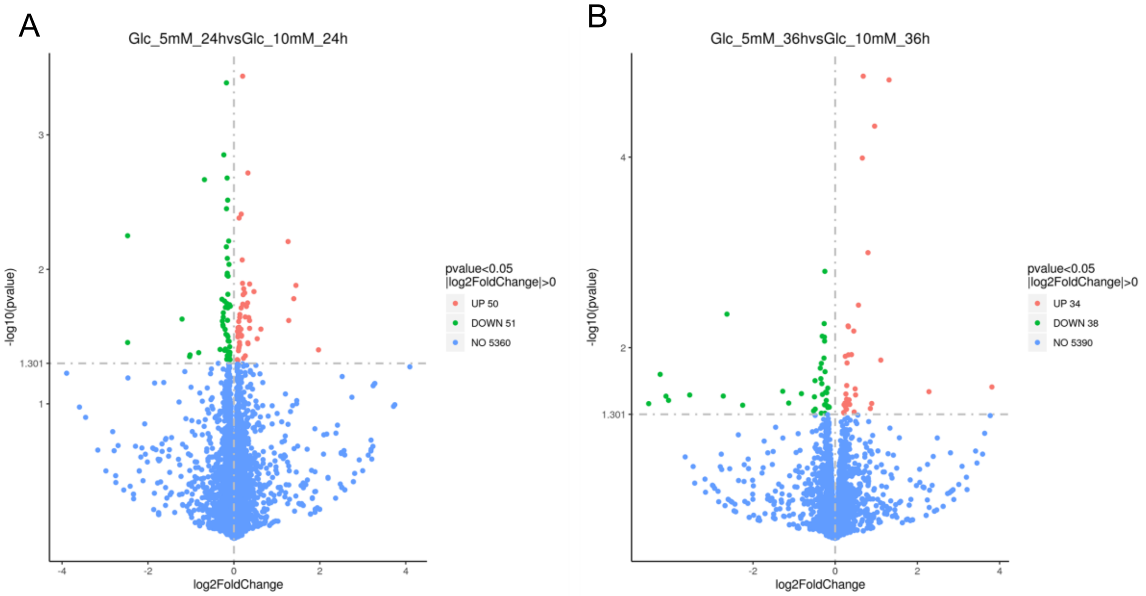


**Figure S1.** Transcriptome analysis of the parasite cultured in different glucose concentrations. Volcano plot showing differential expression of the parasite genes between 5 mM glucose and 10 mM glucose culture conditions at 24 (A) and 36 (B) h.

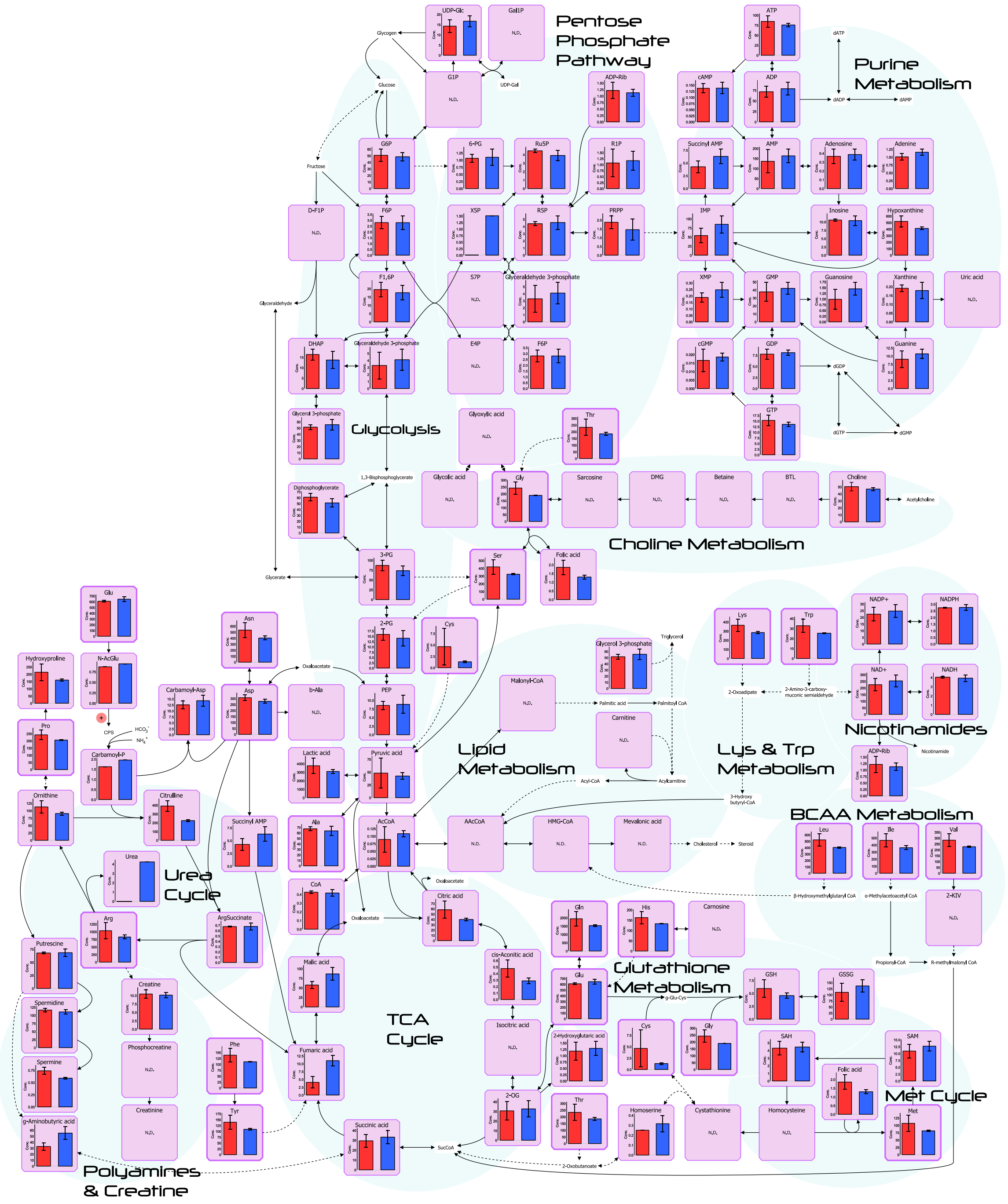


**Figure S2.** Long-term low glucose culture did not change the metabolic profiles of the parasite. The PFK9-mNG expressing parasite was cultured in 5 (red) or 10 (blue) mM glucose for 78–80 h, and 116 metabolites were measured in triplicate.

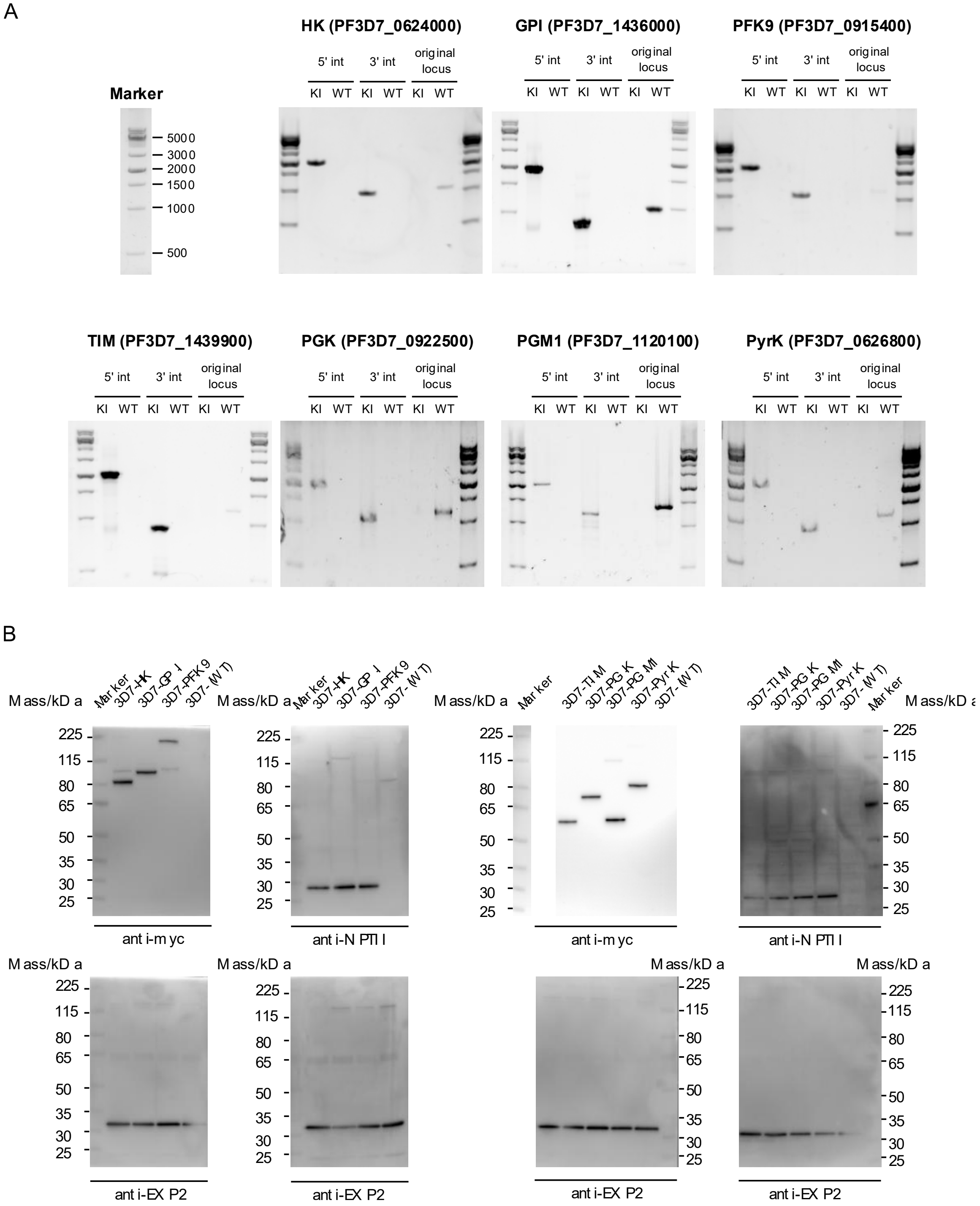


**Figure S3.** Generation of parasites expressing tagged glycolytic enzymes. (A) The results of diagnostic PCR for wild type (WT) and knock-in (KI) strains with different primer sets. (B) C-Terminal tagging of glycolytic enzymes in the *Pf* 3D7 parasite was confirmed by anti-Myc and anti-NPTII western blotting. A total of 5 × $10^6$ parasite lysate per lane was applied. Estimated protein size: HK-mNG-my, 89.4 kDa; GPI-mNG-myc,101.5 kDa; PFK9-mNG-myc,193.6 kDa; TIM-mNG-myc, 62.0 kDa; PGK-mNG-myc, 79.5 kDa; PGM1-mNG-myc, 62.9 kDa; PyrK-mNG-myc, 89.8 kDa; NPTII, 29 kDa; and EXP2, 33.4 kDa.

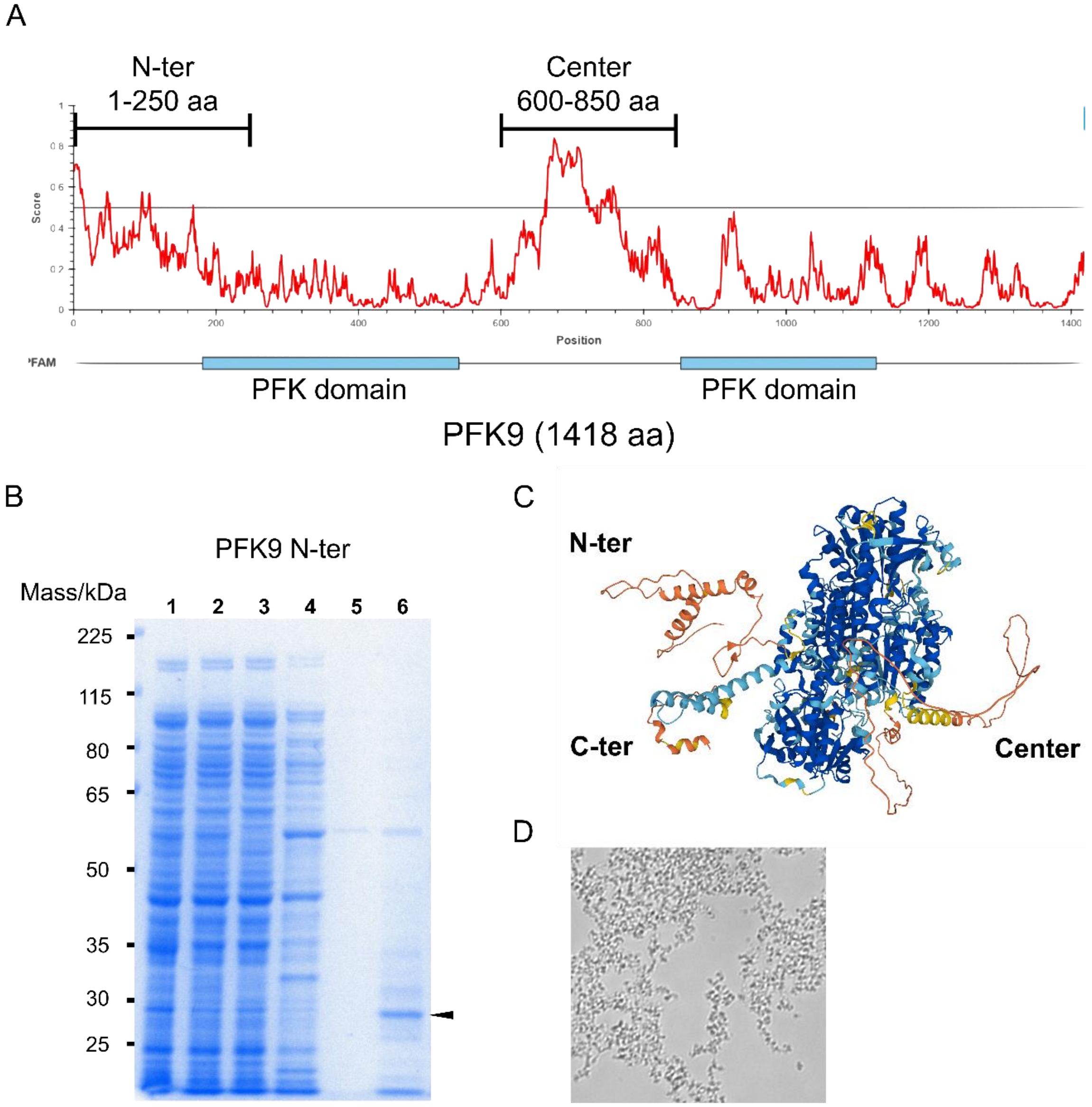


**Figure S4.** Purification of PFK9 intrinsically disordered regions. (A) The result of PFK9 disordered region prediction with IUPred2A and designs of PFK *N*-terminal (*N*-ter) and PFK9 center proteins. (B) The results of SDS-PAGE for PFK9 *N-ter* purification. The arrowheads show the purified target proteins. Lane 1: cell lysate, Lane 2: cytosol, Lane 3: flow through, Lane 4: wash with imidazole-containing buffer, Lane 5: wash with cleavage buffer, Lane 6: purified protein after cleavage. (C) Alphafold2 structure prediction of PFK9. (D) Representative brightfield picture of purified PFK9 *N*-ter protein aggregates.

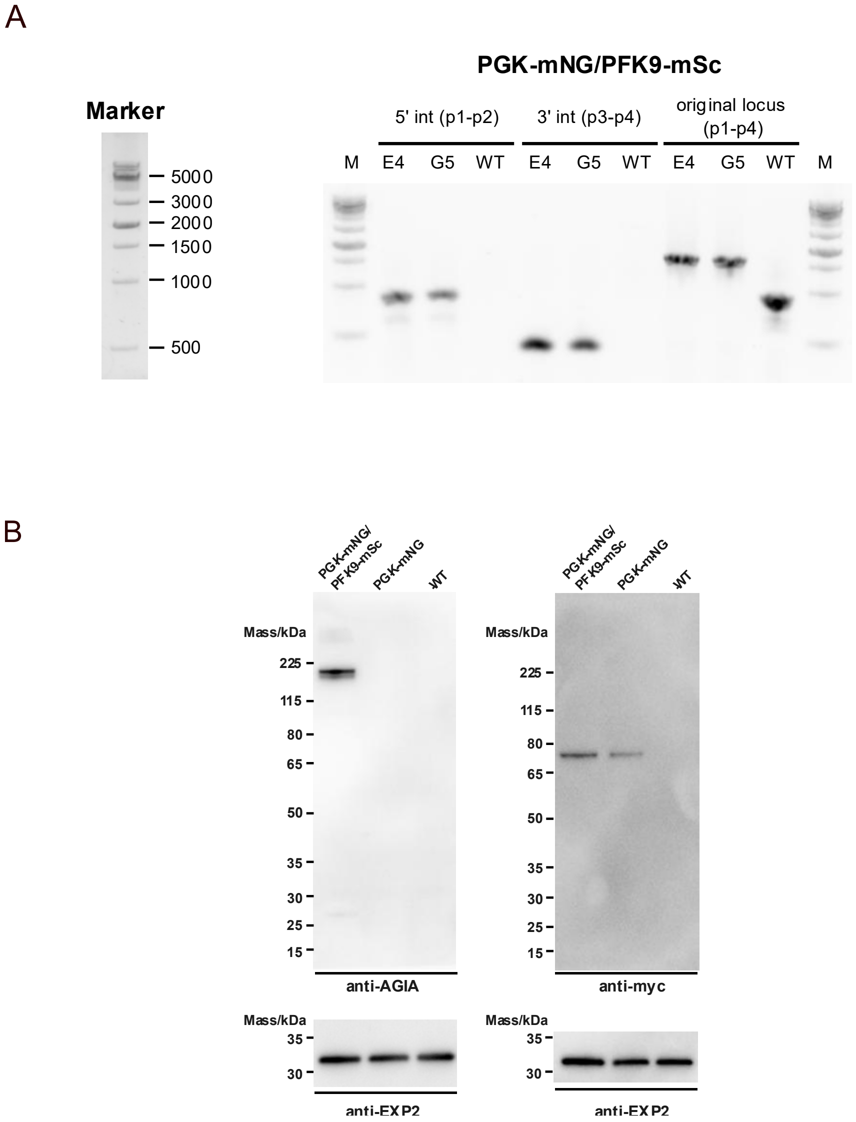


**Figure S5.** Generation of the PGK-mNG/PFK9-mSc double knock-in parasite. (A) The results of diagnostic PCR for the PFK9 locus of wild type (WT) and two double knock-in clones (E4 and G5) with different primer sets. The primer positions are shown in Fig. 5A. M, marker. (B) C-Terminal tagging of PFK9 with mSc and of PGK with mNG in the *Pf* 3D7 parasite was confirmed by anti-AGIA and anti-Myc western blotting. A total of 5 × $10^6$ parasite lysate per lane was applied. Anti-EXP2 was used as a loading control. Estimated protein size: PFK9-mSc-AGIA, 191.4 kDa; PGK-mNG-myc, 79.5 kDa; and EXP2, 33.4 kDa.

**Table S2.** Synthesized DNA sequences used for plasmid construction.

| Name | Sequence |
| --- | --- |
| PfPGK | ACGTAACAGACTTAGGAGGGACCACTACTAGCCATTTTAGGTGGTGCTAAAGTATCAGATAAA<br>ATTCAGTTAATCAAAAATTTATTAGATAAAGTTGATCGTATGATTATTGGAGGTGGTATGGCTTA<br>TACTTTTAAAAAAGTCTTAAACAATATGAAAATTGGTACCTCTCTTTTTGATGAAGCAGGTAGTA<br>AAATTGTTGGAGAAATTATGGAAAAGGCAAAAGCTAAAAATGTTCAAATATTTTTACCTGTAGAT<br>TTTAAAATTGCAGATAATTTTGATAATAATGCTAATACCAAATTCGTTACTGATGAAGAAGGTAT<br>CCCAGATAATTGGATGGGTCTTGATGCTGGTCCAAAAAGTATTGAAAATTATAAAGATGTTATT<br>TTAACATCAAAAACTGTTATTTGGAACGGACCACAAGGTGTTTTCGAAATGCCAAACTTTGCAA<br>AAGGTAGTATCGAATGTCTCAATTTAGTTGTTGAAGTTACCAAAAAAGGAGCTATCACCATTGT<br>TGGAGGTGGAGATACAGCTTCATTAGTTGAACAACAAAATAAGAAAAACGAAATTAGTCATGTA<br>TCTACAGGTGGAGGAGCCTCACTTGAACTTTTAGAAGGAAAAGAATTACCAGGTGTATTAGCA<br>CTTTCAAACAAAATGGTGAGTAAGGGAGAAG |
| PfTIM | ACGTAACAGACTTAGGAGGTAGGACGTTGTTGTTTTTCCTGTTTCCGTACATTATGATCATACA<br>AGGAAATTACTTCAGAGTAAGTTTTCTACTGGTATTCAGAATGTATCAAAATTCGGAAATGGAT<br>CATACACAGGTGAAGTAAGTGCAGAAATTGCCAAGGATTTAAATATTGAATATGTTATTATTGG<br>TCATTTTGAAAGAAGAAAATATTTCCATGAAACCGATGAAGATGTTCGTGAAAAATTACAAGCT<br>TCATTAAAAAATAATTTAAAAGCCGTTGTATGTTTTGGTGAATCTTTAGAACAAAGAGAACAAAA<br>TAAAACTATCGAAGTTATTACAAAACAAGTTAAAGCATTTGTTGATTTAATTGATAATTTTGATAA<br>TGTTATTTTGGCTTATGAACCTTTATGGGCTATTGGTACTGGTAAAACAGCTACACCTGAACAA<br>GCTCAATTAGTACACAAAGAAATCAGAAAAATTGTAAAAGATACATGCGGAGAAAAACAAGCTA<br>ACCAAATAAGAATATTATATGGAGGTAGTGTTAATACTGAAAACTGCTCTTCATTAATTCAACAA<br>GAAGATATTGATGGTTTCTTAGTTGGAAATGCTTCCTTAAAAGAATCTTTTGTTGATATAATAAA<br>AAGTGCTATGATGGTGAGTAAGGGAGAAG |
| PfGPI | ACGTAACAGACTTAGGAGGTAGCAAAATTTATTAAAATTTTCAGCACATATTCAACAATTATCTA<br>TGGAAAGTAATGGAAAATCAGTAGATAGAAATAATCAACCAATCCATTATAATACTTGTCAAGT<br>ATATTTTGGTGAACCTGGTACAAATGGTCAGCATAGTTTTTATCAATTAATACATCAAGGACAA<br>GTTATACCTGTTGAATTAATTGGATTTAAACATTCTCATTTCCCAATTAAATTTGATAAAGAAGTA<br>GTAAGTAATCATGATGAATTAATGACTAACTTTTTTGCACAAGCTGATGCATTAGCTATTGGGA<br>AAACATACGAACAAGTAAAAGAAGAAAATGAAAAAAATAAAATGTCTCCAGAATTATTAACTCAT<br>AAAGTTTTTAACGGTAATAGACCATCAACCTTATTATTATTTGATGAATTAAATTTCTATACATGT<br>GGTTTATTATTATCCTTATATGAAAGTAGAATTGTCGCTGAAGGATTTCTATTAAATATTAACAG<br>CTTTGATCAATGGGGGGTAGAGTTGGGTAAGGTTCTAGCAAAAGAAGTAAGAAATTATTTTAAT<br>GACACAAGAAATCAAAAGAAATCAGATAATACCTATAATTTTAATGAATCTACAAAAATTTTATT<br>AAATTATTACTTGTCCAAAATGGTGAGTAAGGGAGAAG |
| PfPyrK | ACGTAACAGACTTAGGAGGTAGCCTTCCATTTATAGATGCAATTTACAAGGTAAACCAATTATC<br>ACTGCCACTCAAATGTTAGAATCCATGACCAAGAACCCAAGACCAACAAGAGCAGAAGTTACA<br>GATGTAGCTAATGCTGTTTTAGATGGTACTGATTGTGTTATGCTTTCAGGAGAAACTGCAGGTG<br>GTAAATTTCCAGTGGAAGCTGTTACCATTATGTCCAAAATCTGTTTAGAAGCAGAAGCATGCAT<br>TGATTATAAATTATTATATCAATCATTAGTAAATGCAATTGAAACACCAATTAGTGTTCAAGAAG<br>CTGTGGCTAGATCAGCTGTAGAAACAGCAGAATCTATTCAAGCTTCTTTAATTATAGCTTTAAC<br>AGAAACAGGTTATACGGCTAGATTAATTGCCAAATATAAACCAAGTTGTACCATCTTAGCCCTT<br>AGTGCTTCTGATTCAACTGTTAAATGTTTGAACGTACACAGAGGTGTTACATGCATTAAAGTAG<br>GTTCATTCCAAGGAACAGATATTGTTATTAGAAATGCTATTGAAATCGCAAAACAAAGAAATAT<br>GGCAAAAGTTGGTGATAGCGTTATTGCTATTCACGGAATTAAAGAAGAAGTATCTGGAGGTAC<br>CAACTTAATGAAAGTTGTACAAATTGAAATGGTGAGTAAGGGAGAAG |
| PFK9_HR1 | TCAGAATTTGATTGTAAGTTAGCATATGCATATGGACATGCAGCATCAATAGTTATTGAAAGTG<br>GCTTAACAGGATATATTGTATCTATAAGAGGATTATGTGGTAATGTTAAAGATTGGAAATTATTT<br>GCTATACCATTCATATCATTAATGAAAATATTACCAAAAGGACAAGGAAGTAAATATTTAAAAAG<br>TGCATCTAAAGGTGATTTACCAGTAATTCCTAGTGCACCTGTAGATTTGAATGGAAAAGCTTAT<br>AGAAGCTTAAAAATTGCTTTACAAAAATGGCAAATGGAAGATCGTTTTTGTAATCCAGGTCCTA<br>TTCAATTTGAAGGTAATGCATCTAATTATTATAATAGGATATTATTTGAAGAACAATCTGAATATT<br>TTGAAATGTTAAGATATGTTGAATGTTATGCAAATATTTTAAAAGATACTTGTCGTTTTGGTGTC<br>TCAGCTGATTATCTGAAAAACGTTTTTGTTCAGTTGTGTGGTATGTTGGTATTAGCATATAAGCC<br>TAATGATATCTTATCTAATATGCCATATATTGGAAGTATAGAAGATTATTATGATTGGGAAAACC<br>AGAGAAAAAGAATGAACATGGTAAGTAAAGGTGAAGCTGTAATTAAAGA |

| | |
|---|---|
| mScarlet-AGIA | ATGGTAAGTAAAGGTGAAGCTGTAATTAAAGAATTTATGAGATTTAAAGTACATATGGAAGGTA<br>GTATGAATGGTCATGAATTTGAAATTGAAGGTGAAGGAGAAGGTAGACCTTATGAAGGTACAC<br>AAACAGCTAAATTAAAAGTAACAAAAGGTGGTCCATTACCATTTAGTTGGGATATTTTAAGTCC<br>ACAATTTATGTATGGTAGTAGAGCTTTTACAAAACATCCTGCTGATATTCCTGATTATTATAAAC<br>AAAGTTTTCCTGAAGGTTTTAAATGGGAAAGAGTAATGAATTTTGAAGATGGTGGTGCTGTAAC<br>AGTAACTCAAGATACAAGTTTGGAAGATGGTACATTGATTTATAAAGTAAAATTAAGAGGTACA<br>AATTTTCCACCTGATGGTCCTGTAATGCAAAAAAAAACAATGGGTTGGGAAGCTAGTACAGAA<br>AGATTATATCCTGAAGATGGTGTATTAAAAGGTGATATTAAAATGGCTTTAAGATTAAAAGATG<br>GTGGTAGATATTTAGCTGACTTTAAGACTACATATAAAGCTAAAAAACCTGTACAGATGCCTGG<br>TGCTTATAATGTAGACCGTAAATTAGACATAACAAGTCATAATGAAGATTACACAGTAGTAGAA<br>CAATATGAAAGAAGTGAGGGTAGACATAGTACTGGTGGTATGGATGAATTATATAAAGGGCCC<br>GAAGAAGCTGCTGGTATTGCTAGACCATTATAA |

**Table S3.** Primers used for plasmid construction and diagnostic PCR.

| Primer name | Sequence |
|---|---|
| PFK9_F1 | acgtaacagacttaggaggGAGTTTAGCATTATTAAAGAC |
| PFK9_R1 | taattcccggggtgttcatGTTCATTCTTTTTCTCTGGTT |
| HK_F1 | ACGTAACAGACTTAGGAGGGTGCATATCAAAAAGGTAGAGGTAC |
| HK_R1 | TAATTCCCGGGGTGTTCATTGGTAATTGTGGAATGTCC |
| PGM1_F2 | acgtaacagactTAGgaggAGAATTTCAAATTCGACGTTG |
| PGM1_R2 | CTTCTCCCTTACTCACCATTTTTGCTTTTCCTTGGTTG |
| PFK9-mSc_gRNA1_1 | TATTAATGTATTTGTACAGTTATG |
| PFK9-mSc_gRNA1_2 | AAACCATAACTGTACAAATACATT |
| PFK9-mSc-HR1_amp_F | gaattcatatcgataactcTCAGAATTTGATTGTAAGTTAGCATATGC |
| PFK9-mSc-HR1_amp_R | TCTTTAATTACAGCTTCACCTTTACTTAC |
| mSc-AGIA_amp_F | ATGGTAAGTAAAGGTGAAGCTG |
| mSc-AGIA_amp_R | TTATAATGGTCTAGCAATACCAGC |
| PFK9-mSc-HR2_F | AGCTGCTGGTATTGCTAGACCATTATAATGTATATGTTTATGTATT TATCCATGTGTG |
| PFK9-mSc-HR2_R | ccagcctaggagttccatgCCCTTCCAATCAATAGTTTAAAAATATTATAAT GG |
| p2 | CGGTACCTTGACCTACCATATC |
| p3 | caggaaacagctatgaccatg |
| PFK9_p1 | GCTGATGTTGGAAAAAATTATG |
| PFK9_p4 | GAAATATACACACATGGATAAATAC |
| HK_p1 | TTCCCTTGTACATCACCTTCAAT |
| HK_p4 | TCTTCACACACTCACATACAGACAA |
| PGM1_p1 | GTGAAAGTACATGGAACAAGG |
| PGM1_p4 | TATCCACAAATGGGAAACCTCG |
| PyrK_p1 | ACCGCATCAGCTATTAGTTCC |
| PyrK_p4 | ATACATTAAAGAAAGGCAAAGG |
| PGK_p1 | TATTGAAGAAGAAGGTAAAGGTGTTG |

| | |
|---|---|
| PGK_p4 | CATACATACACACATACAATAAG |
| TIM_p1 | AGCAAACTGGAAATGTAATGGAAC |
| TIM_p4 | ATATAGAATACGTTCACCTC |
| GPI_p1 | TTTGGGATTGGGTAGGAGGTC |
| GPI_p4 | AACCCATATTTGCACTTTCACAC |
| Seq.mSc_F | CAAGTTTGGAAGATGGTACATTG |

The lowercase letters indicate the sequences of the plasmid backbone

**Table S4.** Antibodies and reaction conditions used for western blotting.

| Antibody name | Supplier | Dilution factor | Incubation time and temperature |
|---|---|---|---|
| Myc -Tag (9B11) Mouse mAb #2276 | Cell Signaling Technology, Danvaese, MA, USA | 500 or 1000 | 1 hour, room temperature |
| Neomycin phosphotransferase II Antibody (4B4D1) | Novus Biologicals, Centennial, West Grove, CO, USA | 500 or 1000 | 1 hour, room temperature |
| Anti-AGIA Rabbit polyclonal antibody | Ehime University, Ehime, Japan | 1000 | Overnight, 4 °C |
| Anti-EXP2 Rabbit polyclonal antibody | Ehime University, Ehime, Japan | 2500 | 1 hour, room temperature |
| Peroxidase AffiniPure Goat Anti-Mouse IgG (H+L) | Jackson ImmunoResearch, PA, USA | 5000 or 10000 | 30 min, room temperature |
| Peroxidase AffiniPure Goat Anti-Rabbit IgG (H+L) | Jackson ImmunoResearch, USA | 5000 or 10000 | 30 min, room temperature |
